# Seasonal neurogenomic changes provide genetic links between male song and testosterone-mediated neurogenesis in a wild songbird

**DOI:** 10.1101/2025.09.03.674109

**Authors:** Maria E. Mendiwelso, Timothy A. Liddle, Nora H. Prior, Anna K. Magnaterra, Chelsea Haakenson, Katherine A. Stennette, Catalina Palacios, Cara A. Krieg, Gregory F. Ball, Tyler J. Stevenson, Karan J. Odom

## Abstract

Seasonal changes in testosterone mediate the transition from reduced singing in the non-breeding season to high song rates in the early breeding season in male temperate songbirds, a process accompanied by marked neurogenesis in song control regions of the brain. However, the resulting genetic changes and their association with the subsequent behavioral, neuroanatomical and reproductive changes remain poorly understood. Here, we compared gene expression in HVC, a major song control nucleus, between the non-breeding and breeding season in male northern house wrens (*Troglodytes aedon*) and we examined associations with HVC volume, plasma testosterone concentrations, and testes size. Differential gene expression analysis identified three genes (CLIP4, FAM169A, and TTR) consistently linked to seasonal transitions from non-breeding to three breeding stages (pre-laying, egg-laying and incubation). Notably, TTR, which transports thyroid hormone (TH), was highly expressed in the nonbreeding season, consistent with a possible role of TH in regulating seasonal shifts in song output and structure. We also identified seasonal changes in networks of genes related to neural connectivity, cellular restructuring, and cell migration. Weighted gene co-expression network analysis (WGCNA) revealed gene clusters specifically correlated with testosterone and HVC volume. Testosterone-associated genes included genes involved in neural circuit remodeling, chromatin organization, and cytoskeletal dynamics, consistent with testosterone-mediated regulation of these neuroanatomical changes. Genes linked to seasonal increases in HVC volume were involved in neuronal restructuring and neuron migration, implicating these genes in seasonal neurogenesis. Together, our findings link novel gene expression patterns to hormone regulation and neurogenesis underlying seasonal transitions in birdsong.

## Introduction

In temperate songbirds, males exhibit a dramatic annual shift from reduced song rates and relaxed song structure in the non-breeding season to greatly increased rates of stereotyped, complex songs early in the breeding season (Brenowitz, 2004; Catchpole, 2008; Thompson et al., 2007, 2012). These behavioral shifts correspond with seasonal neurogenesis in the song control system (SCS), a specialized neuronal network responsible for song production and modulation in songbirds (Passeri; Nottebohm et al., 1976). Central to this network is the HVC (proper name), a nucleus critical for coordinating song stereotypy and complexity (Alward et al., 2014, 2017; Brenowitz et al., 2015). Because of its seasonal neurogenesis and role in regulating birdsong structure, HVC plays a prominent role in seasonal changes in vocal behavior (Brenowitz et al., 2002; Chen et al., 2013; Ko et al., 2021). Thus, measuring HVC gene expression between the breeding and non-breeding season offers a powerful tool to examine genetic mechanisms that regulate seasonal variation in birdsong.

Photoperiod is the primary environmental cue regulating seasonal changes in physiology and reproductive behavior, including song (Liddle et al., 2022). Increasing photoperiod promotes gonadal growth and secretion of testosterone into blood plasma (Ball, 1999; Ball et al., 2004a; Bernard et al., 1997; Brenowitz, 2004; Watts, 2020). Elevated testosterone promotes neurogenesis and subsequent increases in SCS nuclei volumes, especially in the HVC, leading to increased neuronal recruitment, larger song nuclei volumes, and enhanced neural connectivity (Brenowitz et al., 2015; Tramontin et al., 1999; Thompson et al., 2007; Alward et al., 2017; Chen et al., 2013). These neuroanatomical changes precede peak singing behavior, suggesting they prepare the brain for the demands of reproductive vocal performance (Alward et al., 2013; J. Orije et al., 2020). For instance, male canaries produce a longer, more complex, stereotyped song during the breeding season when testosterone concentrations are highest but revert to more variable songs as testosterone declines (Alward et al., 2014; Nottebohm et al., 1987). Testosterone in HVC is specifically known to promote increased syllable usage and sequencing, leading to more complex songs in the breeding season (Alward et al., 2017). This photoperiodic regulation of hormones and breeding behavior is a widespread mechanism, governing gonadal growth and the onset of reproduction at large in many vertebrates (Ball, 1999; Ball et al., 2004b).

While we have a solid understanding of the neuroendocrine basis of seasonal change in song, we are only beginning to examine the underlying genetic mechanisms via large-scale changes in gene expression (e.g., transcriptomic analyses; see Clayton, 2013;Orije et al., 2022). Specifically, few studies have simultaneously examined seasonal variation in bulk transcriptomic changes in gene expression within the HVC alongside changes in neuroanatomy, hormone concentrations, and singing behavior – much less in wild songbirds. Transcriptional changes in the HVC may represent early molecular events initiating seasonal remodeling of this nucleus, ultimately giving rise to downstream hormonal, anatomical, and behavioral modifications (Lovell et al., 2018; Thompson et al., 2008). There is substantial variation across songbird species in the hormonal regulation and seasonal timing of song and reproductive behavior; however, most research on this topic has only focused on a few captive species (Goymann et al., 2014; MacDougall-Shackleton et al., 2009; Odom et al., 2025). This natural variation likely underscores differences in the regulatory mechanisms across species (Brenowitz, 2004; Goymann et al., 2014). Therefore, identifying the gene expression patterns that precede or coincide with HVC structural and functional changes in different songbird species can shed light on the molecular drivers of plasticity and the molecular processes underlying evolutionary transitions in seasonal reproductive behavior, including song (Colquitt et al., 2023; Rasika et al., 1994).

Emerging evidence suggests that testosterone may interact with other endocrine signals— such as thyroid hormones (TH)— to regulate seasonal plasticity in birdsong (Orije et al., 2022). Moreover, discrepancies in the timing between testosterone peaks and the onset of neurogenesis or increased singing suggest a more complex regulatory network, where gene expression changes may precede or mediate downstream hormonal and neural effects (Caro et al., 2005; Tramontin et al., 2001). Examining changes in gene expression is one way to elucidate the mechanisms involved in this intricate set of seasonal changes.

To address this gap, we investigated the seasonal variation in gene expression within the HVC and its association with changes in HVC volume, circulating testosterone, and testes size in naturally breeding male northern house wrens (*Troglodytes aedon*). Using transcriptomic analyses on HVC tissue samples collected across different breeding stages, integrated with precise neuroanatomical measurements and hormone assays, we aim to provide new insights into the molecular basis of seasonal plasticity in the SCS.

## Materials and methods

The northern house wren, an abundant small passerine bird widely distributed across the United States and Canada with nonbreeding ranges into Mexico, provides an excellent model for studying the male SCS. Northern house wrens exhibit clear seasonal changes in singing behavior, with males singing intensely early in the breeding season to attract mates and establish territories (Johnson, 2020). This behavior makes this species well-suited to study the genetic and hormonal underpinnings of song control.

### Field sites, data & sample collection

For this study, we focused exclusively on male northern house wrens because they show dramatic seasonal shifts in singing behavior from little to no song in the non-breeding season to sometimes over 1000 songs by a single male in the first few hours of the morning in the early breeding season (Odom et al. in review). Males were captured at four different times of the year: non-breeding, pre-laying, egg-laying, and incubation. These breeding stages were determined by the precise breeding stage of each male’s mate, which could be readily monitored in these nestbox-breeding birds. Male northern house wrens do not incubate or lay eggs; these stages refer to the breeding stage of their mate. We chose these three breeding stages because our previous research showed that male song rates and testosterone are highest pre-laying and then steadily decrease until the pre-laying stage of the next breeding cycle (See Odom et al. in review, Liddle et al. in review, Cramer, 2012). Non-breeding males (n = 7) were collected in February 2022 at the Hague Dairy Unit, University of Florida, Alachua County, FL. Breeding birds were collected May – early July 2022 and 2023 during the following specific timeframes: pre-laying stage (n = 5, in early May), egg-laying stage (n= 6, mid-May to early June), and incubation stage (n = 6, end of May and early June to early July). Breeding birds were captured on Pennsylvania State Game Lands 127 (Monroe County), 300 (Lackawanna County), and 310 (Wayne County; See supplementary table 1 for a full list of sample dates and locations).

Northern house wrens were captured using mist nets. Most individuals were captured using a brief period (less than 5 minutes) of song playback as an audio lure. Some individuals were captured prior to playback by flushing them into the net. Most individuals were successfully captured and collected within 10 minutes after the initial onset of playback. However, four individuals (two non-breeding, one pre-laying, and one incubating) were captured more than 45 minutes post-playback.

Upon capture, tissue was immediately collected in the field: following rapid decapitation, the whole brain was quickly removed. After removal, brains were quickly bisected on ice using a clean razor blade. The left hemisphere of each brain was immediately flash-frozen on powdered dry ice and then stored at -80°C until RNA extraction and sequencing. The right hemisphere was fixed in 4% formaldehyde for 24 hours, then stored in 30% sucrose and PBS for 1-3 days before being frozen on dry ice and stored at -80°C for histological analysis. Most brains were processed (either flash-frozen or immersed in fixative) within 7-11 minutes post-lethal collection, with a few exceptions taking up to 15 minutes for processing.

Immediately after decapitation, all trunk blood was collected in a 1.5mL microcentrifuge tube coated with 20 uL of heparin to prevent coagulation and stored on ice up to 8 hours before centrifugation at 10,000 rpm for 15 min. The plasma was then pipetted into a clean 0.5 mL microcentrifuge tube and stored at -20°C for the remainder of the field season (up to 2 months) and then at -80°C until assayed (up to 1.5 yr later).

All work was conducted in accordance with the Association for the Study of Animal Behavior (ASAB) guidelines and the Ornithological Council’s Guidelines to the Use of Wild Birds in Research. Our methods were approved by the Institutional Animal Care and Use Committee (IACUC; #FR-APR-21-17), University of Maryland, College Park. These activities were permitted under Federal U.S. Fish and Wildlife Service Scientific Collecting Permit number MB01550B, Pennsylvania State Game and Conservation Commission Scientific Collecting Permit number 52256, Pennsylvania State Game and Conservation Commission Specific Use Permit number 52524, Florida Fish and Wildlife Conservation Commission Scientific Collecting Permit number LSSC-12-00016C, and Virginia Department of Wildlife Resources Scientific Collecting permit number 3391840.

### Reproductive physiology measurements

Morphometric and hormonal data were collected for all males. Each individual was weighed in a bird bag prior to collection using a 30g Pesola scale. The bag weight was then subtracted to calculate the weight of each bird. Soon after collection and immediately after dissection, left testis height and width, and right testis height and width were measured in the field on the fresh tissues using high-precision calipers. Average testes size was calculated as the mean of the left and right values for testis area (mm^2^) height × width for each testis. In addition, body mass, bill length, wing length, and tarsus length were measured for each bird in the field.

### Testosterone assays

We used a Testosterone ELISA kit (Cayman Chemicals #582701) to assay all blood plasma samples. The assay was performed with 20μL of plasma per well for all non-breeding males in singleton and 4μL of plasma for all males in duplicate. Intra-assay variation was calculated based on a set of plasma samples and two reference standards of known testosterone concentration (0.5 and 0.25 ng/mL) run on each plate. Plasma testosterone concentrations were interpolated from raw absorbance as log concentrations data using the RIA & ELISA template in GraphPad Prism 9.5.0 (GraphPad Software, LLC). The assays for 2022 and 2023 were run in separate years within 5 months following each field season on two separate plates each year. For each of the two years, we achieved the following intra- and inter-assay variation: 2022 – intra-assay variation = 7.34, 9.12, inter-assay variation = 15.62; 2023 – intra-assay variation = 7.88, 9.94, inter-assay variation = 13.02.

### Histology & HVC volume calculations

Brain tissue was sectioned in the coronal plane into 4 series 30 μm thick with a cryostat (Microm HM 500 OM). Immunocytochemistry (ICC) was performed to visualize Hu proteins. Hu is a nuclear stain for neurons that can be used to calculate neuron cell counts and cell density. In the current study, we also found that Hu provided a strong contrast for the boundary surrounding HVC, including in non-breeding males, which could be difficult to see with Nissl for some non-breeding individuals. Therefore, for this study we also used Hu to calculate HVC volume. Our ICC assays were run using well established immunohistochemical protocols (Lynch et al., 2012). Briefly, free-floating sections were washed in phosphate-buffered saline (PBS, 0.01 M, pH 7.5) and treated with 0.1% sodium borohydride. After three rinses in PBS, sections were then treated in 0.5% H2O2 for 30 min to block endogenous peroxidases. After another three rinses in PBS containing 3% Triton-X (0.3% PBST), sections were placed for 1.5 h in blocking solution (20% normal horse serum (NHS) in PBST). Sections were then incubated for 48h at 4 °C in 2% NHS and primary antibody (1:4000, Hu mouse monoclonal IgG invitrogen: A21271, Walton et al. 2012) in PBST. Following three rinses in PBS, sections were then incubated for 1 h in secondary antibody (Biotinylated horse anti-mouse, 1:250, Vector, cat#: BA-2000) in 0.1% PBST. Antibody bound to Hu protein was amplified using Vectastain ABC Elite kit (Vector, cat#:PK-6100) and visualized using 3,3-diaminobezidene tetrahydrochloride chromagen DAB, with nickel enhancement for a black reaction product (Vector, cat#: SK-4100). Sections were then mounted on slides and cover-slipped for analysis via microscopy. All slides were photographed using a Zeiss Axioskop light microscope equipped with an Axiocam 305 Color camera, and images were acquired with Zeiss ZEN 3.8 software with default parameters.

To visualize Hu, all sections for each bird that contained HVC were photographed at 5x. All images were saved in a tiff file format to preserve dimension data, labelled without treatment identifying information, and saved to be later processed in Image J. For each image, the boundary of HVC was outlined on the image using the shape drawing tool and the shape was added to the tiff overlay for later reference. This was repeated for every section on which HVC was visible. HVC volumes were then calculated from the outlined areas in ImageJ by multiplying each area by the section thickness (0.03 mm), summing these values, and then multiplying this value by 4, since only every fourth section was stained and measured (Alward et al., 2013, 2016).

### Assessing physiological and neuroanatomical changes between breeding stages

To assess whether these physiological and neuroanatomical measurements changed significantly between the breeding and non-breeding season or across breeding stage, we used an ANOVA with post-hoc Tukey’s tests (p < 0.05) for the testes size, testosterone concentration, and HVC volume.

### Tissue sample preparation for RNA extraction

Within one month prior to RNA extraction, all flash-frozen left hemispheres of northern house wren brains were sectioned and micro-dissected following Palkovit’s punch technique (George and Rosvall 2025, Heimovics et al., 2016, 2012; Palkovits, 1983). To collect punches, brains were first sectioned in the coronal plane at 200 μm on a cryostat at -16 to -18°C. Sections were thaw-mounted on microscope slides and all microdissections were made while in the cryostat at -25 to -20°C. All HVC punches were made as three lateral to medial punches using a 0.5 mm (inner diameter) stainless-steel sample corer (catalogue #18035-50; Fine Science Tools, Foster City, CA, USA). The specific sections and locations to punch were verified using Nissl and Hu-stained tissue from the formaldehyde fixed right hemisphere of each brain. The final punch locations for house wrens were determined based on the distance (number of sections) from major white-tract landmarks including TrSM split, anterior commissure (CoA) and CoA + Tractus occipito-mesencephalicus (OM). Specifically, we punched HVC rostral to caudal across four sections starting two 200 μm sections after first appearance of CoA + OM. This allowed us to capture majority of HVC for both breeding and non-breeding males. All tissue punches were stored in 1.5 mL microcentrifuge tubes at -80°C until RNA extraction.

### RNA extraction

Tissue punches were first homogenized using a handheld tissue homogenizer in 250μl TRIzol reagent in 1.5ml Eppendorf tubes before incubation at room temperature (RT) for 5 minutes. 50μl chloroform was added and tubes were inverted 10x to mix. Tubes were incubated for 3 minutes at RT and then spinning in a centrifuge at 4°C for 15 minutes at 12,000g. Following centrifugation, the aqueous phase containing the RNA was carefully removed by pipetting.

RNA samples were treated to eliminate DNA by adding 20μl Buffer RDD (Qiagen), 5μl DNase I stock solution, and 50μ RNase-free water and incubating at RT for 10 minutes. Then, 700μl Buffer RLT was added and mixed to each tube before adding 500μl 100% ethanol and mixing by pipetting. Each sample was transferred to a RNeasy MinElute spin column (chilled at 4°C beforehand) and centrifuged for 15 seconds at 8,000g, discarding flow through. Columns were washed with 500μl 80% ethanol and centrifuged for 15 seconds at 8,000g, discarding flow through. Columns were placed in 2ml collection tubes and 500μl Buffer RPE was added to the columns. Then these were centrifuged again for 15 seconds at 8,000g before adding 500μl 80% ethanol and centrifuging again in the same manner. Tubes were then centrifuged for 5 minutes at 8,000g to dry the column membranes. Columns were placed in 1.5ml collection tubes and 14μl RNase-free water was added to each column membrane. Tubes were centrifuged for 1 minute at full speed to collect the purified RNA in solution. Extracted RNA concentration and purity was quantified using a NanoDrop spectrophotometer before storing it at -80°C.

### RNA Sequencing and Functional Annotation

RNA was sequenced according to the protocol outlined by the Oxford Nanopore Technologies SQK-PCB111.24 PCR-cDNA sequencing barcoding kit. The resultant cDNA was sequenced using MinION (Oxford Nanopore Technologies). Each sequencing run had a total runtime of 72 hours at a voltage of -180mV.

Fast5 reads were basecalled and demultiplexed with the guppy base caller (Oxford Nanopore Technologies). Adapters were removed from reads, and short reads were filtered out using Porechop and Filtlong, respectively (R. Wick, 2021; R. R. Wick et al., 2017). Only reads of >25bp were kept.

Transcripts were aligned to a reference *Taeniopygia guttata* genome (zebra finch; https://www.ncbi.nlm.nih.gov/datasets/genome/GCF_000151805.1/). Our sequencing analyses produced a transcriptome for the northern house wren HVC with an average N50 of 567 bases, an average number of reads per sample of 452,000, and an average Phred quality score of 21.1, indicating a basecalling error rate of <1%. Transcript expression was quantified using Salmon (Patro et al., 2017) and then imported into edgeR.

### Differential Expression Analysis

Differential gene expression analysis was conducted using edgeR to identify differentially expressed genes (DEGs) (Robinson et al., 2010). Gene counts were normalized using the Trimmed Mean of M-values (TMM) method to account for differences in library sizes. Pairwise comparisons were conducted between the non-breeding stage and each reproductive stage (pre-laying, egg-laying, and incubation) using a quasi-likelihood framework. Hierarchical clustering analysis was performed using Euclidean distance and Ward’s minimum variance method on log-transformed expression data (Ward, 1963) and visualized along DEGs expression using heatmaps. Principal component analysis (PCA) was performed using edgeR to evaluate sample clustering and detect potential batch effects. Transcript-level counts, filtered by DRIMSeq, were normalized using the TMM method and centered (but not scaled) prior to singular value decomposition. Genes were considered significantly differentially expressed if they met the criteria of an adjusted p-value (FDR) < 0.01 and a log2 fold change (logFC) > |1|. Volcano plots were used to visualize up and downregulated genes, and normalized counts of selected genes of interest were plotted to assess expression patterns.

### Weighted Gene Co-Expression Network Analysis

Transcript counts from salmon were imported into DESeq2 (Love et al., 2014) and subjected to variance-stabilizing transformation (VST). The resulting normalized expression matrix was used as input for weighted gene co-expression network analysis (WGCNA) to cluster genes with similar expression patterns and identify functionally relevant gene modules (Langfelder et al., 2008). Network of co-expressed genes and module detection were performed using the one-step approach in WGCNA. Z-scores (|Z| > 3) were calculated to detect gene-specific outliers, and individual I7 did not surpass the threshold (Supplementary Fig.1), and thus this individual was excluded from subsequent analysis. To ensure the robustness of the analysis, particular attention was given to the selection of the soft-thresholding power (β), a key parameter that influences the balance between gene connectivity within modules and the preservation of scale-free topology (R^2^). Increasing β enhances the scale-free topology fit (R^2^) while reducing gene connectivity, leading to fewer modules containing a higher number of genes with lower co-expression strength. For this study, a β value of 4 was chosen to construct the adjacency matrix, achieving a signed scale-free topology fit (R^2^ ≥ 0.8) and an average gene connectivity of 21.1 (Supplementary Fig 2). Alternative β power values (e.g., 3 and 5) were also evaluated, yielding consistent clustering of DEGs into specific co-expression modules. This stability across varying β values reinforces the robustness of the identified gene co-expression patterns. Subsequently, merged modules were created after applying the module eigengene similarity threshold (Supplementary Fig 3).

We further compared the co-expressed gene modules from WGCNA analysis with the results from DEG analysis, to identify which co-expression modules were associated with our candidate DEGs. Additionally, module eigengenes were correlated with phenotypic traits (testosterone concentrations, testes size, HVC volume) using Spearman’s correlation (FDR-adjusted p-values). Significantly correlated modules were further analyzed based on literature search to explore their biological relevance.

## Results

### Reproductive physiology and neuroanatomy

Using ANOVA and post-hoc Tukey’s tests we found that testis size and HVC volume differed between the non-breeding and breeding season, but not among the breeding stages. Average testes size was significantly larger in the breeding season compared to the non-breeding season (F_(3,18)_ = 96.06, p < 0.001; Fig. 2a). Post-hoc Tukey’s tests revealed that testis size was not significantly different among breeding season stages (pre-laying, egg-laying, and incubation; Supplementary Fig. 5). Similarly, HVC volume was significantly larger in the breeding season than the non-breeding season F_(3,17)_ = 6.91, p < 0.001; Fig. 2c) but was not significantly different among the breeding stages (post-hoc Tukey’s tests; Supplementary Fig. 6). In contrast, testosterone concentrations varied significantly across the non-breeding and breeding season and among the breeding stages (F_(3,19)_ = 5.97, p < 0.005; Fig. 2b), with levels significantly higher during the pre-laying stage to all other stages (post-hoc Tukey’s tests; Supplementary Fig. 7).

### Differential Expression Analysis and Gene Categories

We identified differentially expressed genes (DEGs) between each breeding stage when compared to non-breeding birds. In the non-breeding vs pre-laying comparison, 16 out of 2,858 tested genes were differentially expressed, with 10 showing upregulation and 6 showing downregulation (Supplementary Fig. 4a). For the non-breeding vs. egg-laying comparison, 32 out of 3,154 genes were differentially expressed, including 17 upregulated and 16 downregulated genes (Supplementary Fig. 4b). In the non-breeding vs. incubation comparison, 24 out of 2,992 genes exhibited differential expression, with 13 upregulated and 11 downregulated (Supplementary Fig. 4c). Across all comparisons, a total of 58 DEGs were identified (FDR < 0.01 and log2FC > |1|).

There was minimal overlap of identified DEGs among pairwise comparisons —non-breeding vs. pre-laying, non-breeding vs. egg-laying, and non-breeding vs. incubation. Only three DEGs were differentially expressed across all three comparisons: CAP-Gly Domain Containing Linker Protein Family Member 4 (CLIP4), Family With Sequence Similarity 169 Member A (FAM169A), and Transthyretin (TTR) - (Fig. 1a). CLIP4 (Fig. 1c) and FAM169A (Fig. 1d) exhibited a general upregulation from the non-breeding to the breeding stages, whereas TTR showed downregulation when transitioning from the non-breeding to the breeding season (Fig. 1e). Differential expression of CLIP4, FAM169A, and TTR across all stages highlight their potential role in mediating transitions between non-breeding and breeding states.

**Figure. 1.**
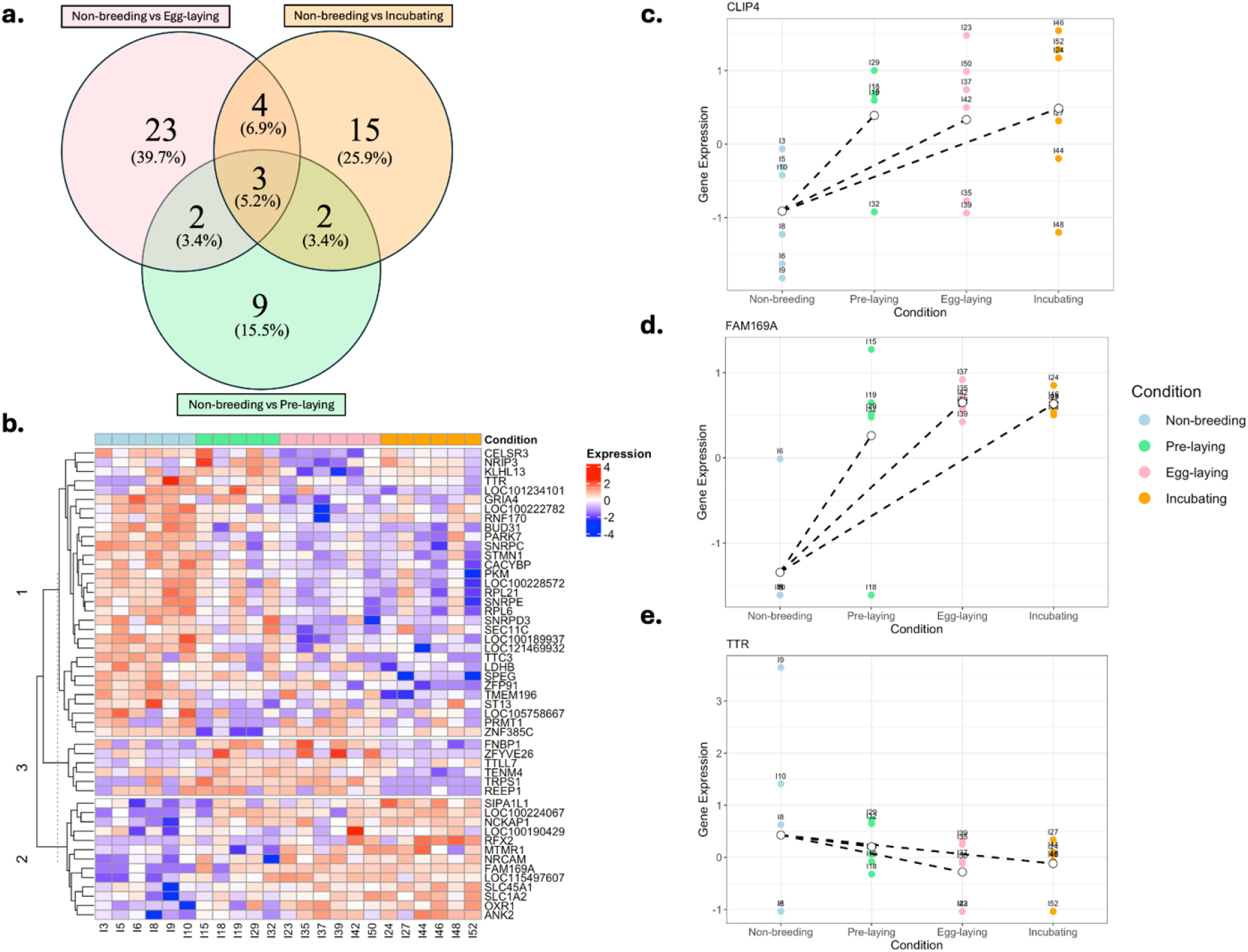
**(a)**. Venn diagram illustrating non-breeding vs. breeding stage comparisons. Most differentially expressed genes DEGs are unique to each stage. Only three genes were commonly differentially expressed across all comparisons. **(b)**. Hierarchical clustering analysis and gene expression of the top 50 DEGs (ranked by false discovery rate, FDR) for each breeding season. Columns are color-coded by season: blue (non-breeding), green (pre-laying), pink (egg-laying), and orange (incubation). **(c-e)** Gene expression profile for the three genes that were commonly differentially expressed across all comparisons **(c)** CLIP4, **(d)** FAM169A, and **(e)** TTR. Colored points represent individual samples. White points represent mean expression by season, with lines connecting non-breeding season with each breeding stage.

Between two comparisons there was also limited overlap: the non-breeding vs. pre-laying and non-breeding vs. incubation comparisons shared five DEGs, as did the non-breeding vs. pre-laying and non-breeding vs. egg-laying comparisons. The non-breeding vs. egg-laying and non-breeding vs. incubation comparisons shared seven genes.

The limited number of shared DEGs among comparisons indicates that most transcriptional changes are stage-specific, meaning that most genes we identified as differentially expressed between the nonbreeding season and each breeding stage were specific to each breeding stage. Hierarchical clustering analysis identified three distinct gene expression clusters (Fig. 1b). Cluster 1 grouped genes that showed high expression during the non-breeding season and low expression during the breeding season. This cluster included TTR—a DEG shared across all comparisons with a well-established role in thyroid hormone transport and neuroplasticity—as well as genes such as CELSR3, Glutamate Ionotropic Receptor AMPA Type Subunit 4 (GRIA4), Ribosomal Protein L6 (RPL6), and Protein Arginine Methyltransferase 1 (PRMT1), broadly associated with synaptic connectivity, excitatory neurotransmission, protein synthesis, and neuronal differentiation. Conversely, genes in cluster 2 showed low expression during the non-breeding season and high expression during the breeding season. Among these genes, Neuronal Cell Adhesion Molecule (NRCAM), Solute Carrier Family 1 Member 2 (SLC1A2), and Ankyrin 2 (ANK2) are broadly associated with synaptic function and neuronal connectivity. Other stage-specific DEGs within this cluster included FAM169A and Oxidation Resistance 1 (OXR1). Finally, cluster 3 exhibited low expression in the non-breeding and incubation stages, and high expression during the pre-laying and egg-laying stages. Notably, this cluster included Tubulin Tyrosine Ligase Like 7 (TTLL7), a gene essential for post-translational tubulin modification, and Teneurin Transmembrane Protein 4 (TENM4), linked to axon guidance and neuronal connectivity, as well as stage-specific DEGs such as Formin Binding Protein 1 (FNBP1).

### Weighted Gene Co-Expression Network Analysis (WGCNA) and correlation with phenotypic traits

WGCNA identified 18 modules of co-expressed genes that included 3,730 transcripts (Fig. 3a.). Only 296 genes were not included in any co-expression module (Module 0). Modules 1, 2, 3, and 10 collectively encompass more than half of the total number of DEGs (n = 36; Fig. 3a; Supplementary Table 2). Among these, modules 1 (19 DEGs) and 10 (7 DEGs) exhibited the highest percentage of DEGs relative to the total number of genes in each module (Fig. 3a).

**Figure 2.**
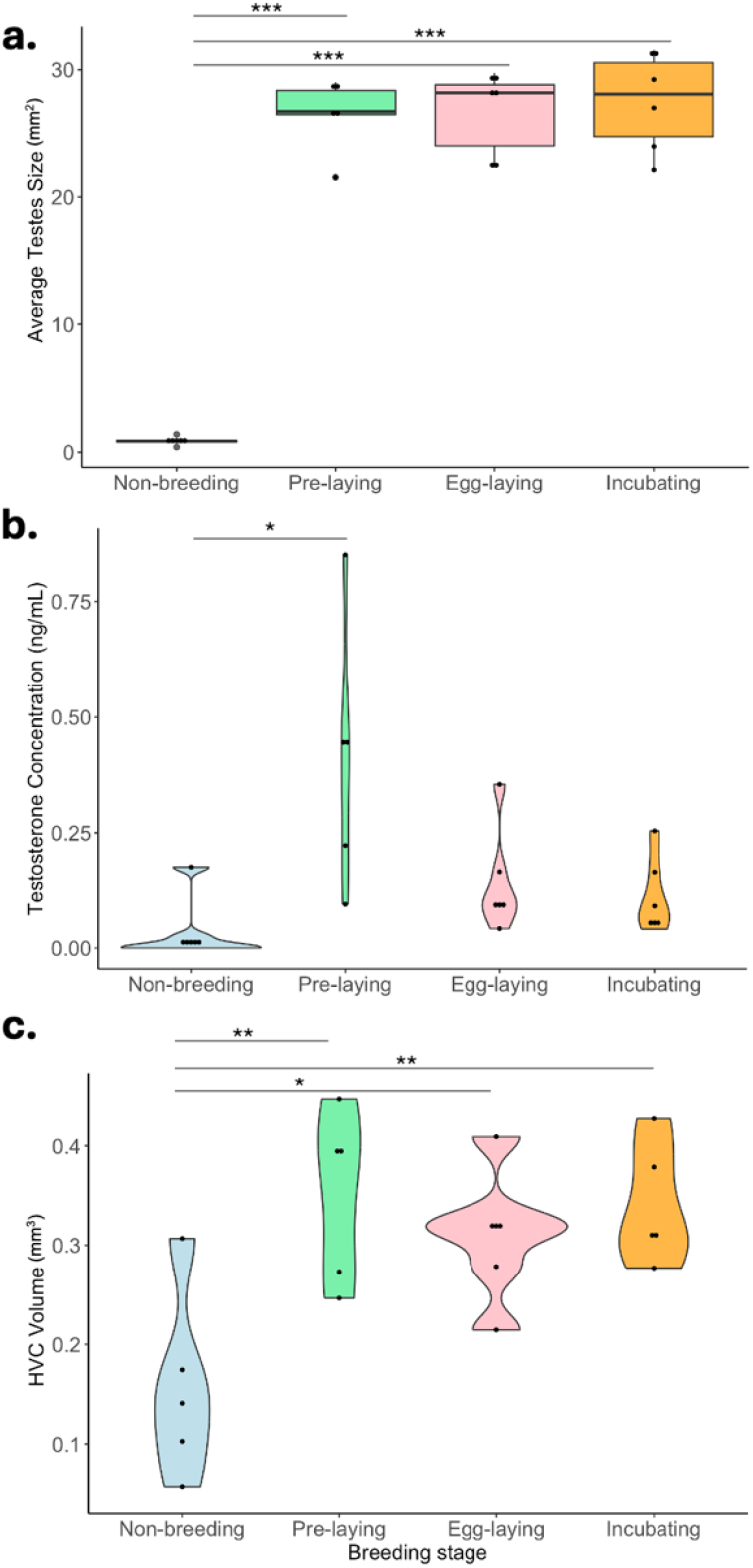
**(a)** Average testes size (mm^2^) (**b)** testosterone concentration (ng/mL) **(c)** HVC volume (mm^3^) across four reproductive stages: non-breeding, pre-laying, egg-laying, and incubating. ANOVA revealed a significant effect of breeding stage on testosterone concentration (F =5.97 *p* = 0.00478), testes size (F = 96.06, p < 0.001), and HVC volume (F = 6.91, p< 0.01). Significance levels: * p < 0.05; ** p < 0.01; *** p < 0.001

**Figure 3.**
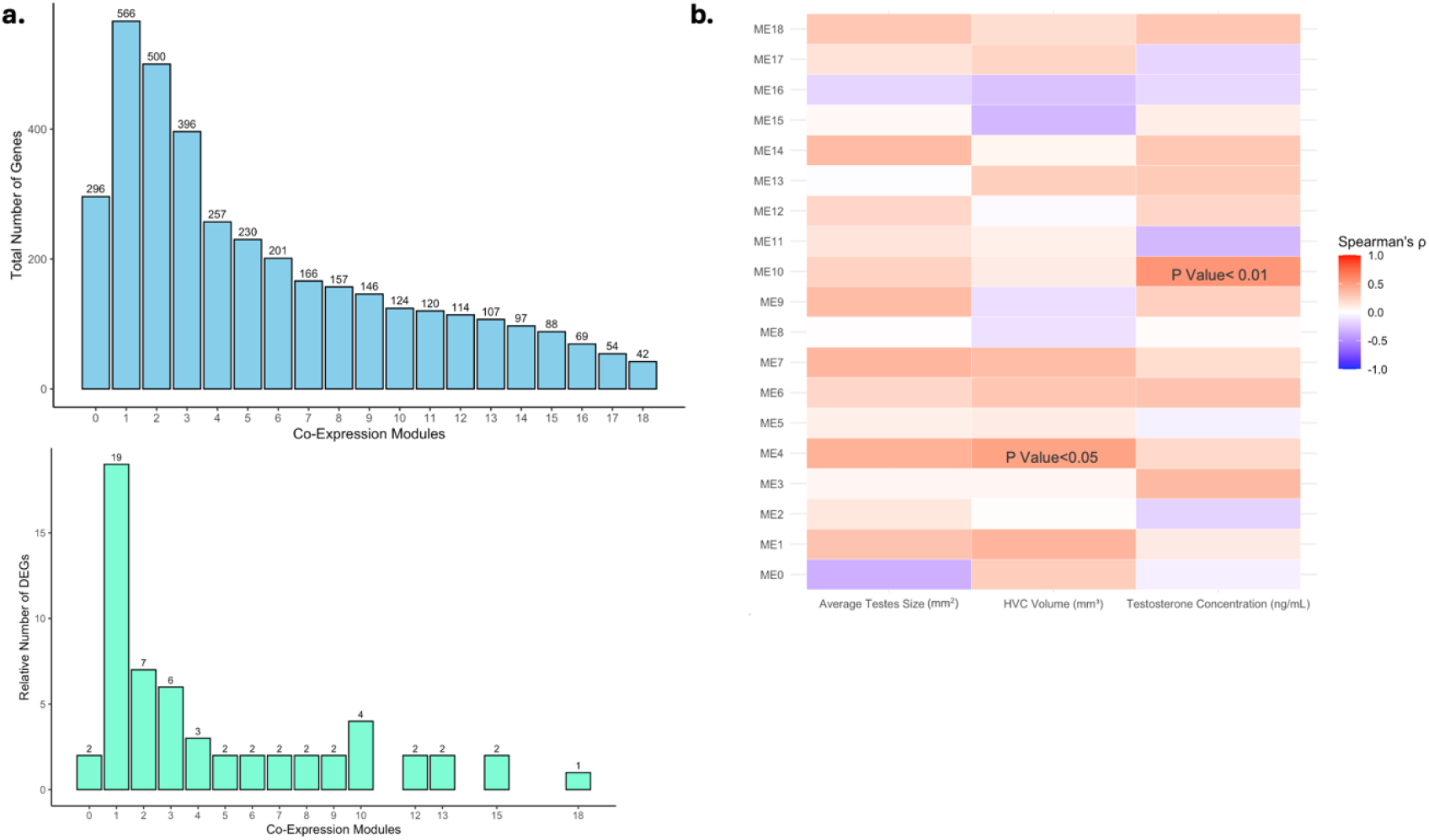
**(a)**. Upper panel bar plot shows the total number of genes assigned to each co-expression module based on WGCNA analysis. Genes not assigned to any module are shown as module 0. Lower panel bar plot shows the total number of DEGs in each module; the total number of DEGs is shown on top of the bars. ***(b)*** Heatmap displaying Spearman’s correlation coefficients (ρ) between co-expression modules and examined traits: average testes size (mm^2^), testosterone concentration (ng/mL) and HVC Volume (mm^3^). The color gradient indicates the magnitude and direction of the correlation: red tones represent positive correlations, while blue tones indicate negative correlations. For significant correlations, the corresponding p-values are shown within the heatmap.

Module 10 was the most strongly, positively correlated with testosterone concentration (Spearman’s ρ = 0.54, p = 0.0075). DEGs in this module included Cytoplasmic Linker Associated Protein 1 (CLASP1), Protein Phosphatase 1 Regulatory Subunit 12B (PPP1R12B), Transmembrane Protein 196 (TMEM196), and FAM169A. FAM169A is one of the DEGs that was upregulated in all three breeding stages compared to the non-breeding season. We noted that module 3 contained TTR, a gene of interest due to its established role in thyroid hormone transport. Although the correlation between module 3 and testosterone concentration was not statistically significant (ρ = 0.36, p = 0.0877), the differential expression of this gene between non-breeding and breeding seasons, along with the known interplay between thyroid hormones and reproductive physiology, suggests that TTR and other genes in this module may warrant further investigation.

For HVC volume, module 4 showed the strongest positive correlation (ρ = 0.48, p = 0.0309), suggesting a potential link between the expression of genes in this module and neural plasticity, including seasonal SCS development. DEGs within this module included LOC115497607, TTLL7 and Tousled Like Kinase 1 (TLK1; Fig. 3b).

No significant correlations were observed between co-expression modules and average testes size. Due to the limited number of DEGs within the modules, a Network Functional Enrichment Analysis was not feasible. Instead, we conducted a comprehensive literature review of the DEGs to infer potential functional associations.

## Discussion

Our results reveal associations between seasonal changes in gene expression and shifts in both physiological and phenotypic traits. Specifically, we identified key DEGs linked to reproductive transitions, including CLIP4, FAM169A, and TTR, which were consistently differentially expressed in all three comparisons between non-breeding and breeding stages (pre-laying, egg-laying, and incubation). Most DEGs exhibited stage-specific regulation when comparing non-breeding birds to individual breeding stages. However, the presence of shared DEGs in HVC between non-breeding and all three breeding stages suggests a core involvement of these genes in seasonal change in song. Using WGCNA, we identified several genes, such as FAM169A, CLASP1, PPP1R12B, and TMEM196, that showed strong associations with circulating testosterone concentration, suggesting a potential role of these genes in hormone-driven modulation of seasonal singing behavior. Expression levels of TTLL7 and TLK1 were specifically linked to variation in HVC volume, implicating a function of these genes in seasonal neurogenesis within this song control nucleus. These findings highlight the dynamic interplay between gene expression, hormone concentrations, and neural plasticity in shaping seasonal reproductive transitions.

### Breeding stage-specific genes

The stage-specific nature of most DEGs indicates that the HVC undergoes dynamic and temporally precise transcriptional shifts across the reproductive cycle. Rather than reflecting a global breeding-related adjustment, our results suggest that distinct molecular modules are recruited at each stage, consistent with the idea that different aspects of reproductive behavior and neural plasticity are orchestrated by distinct gene expression (Lovell et al., 2008; Mello et al., 2004).

One group of genes showed high expression in the nonbreeding period and a consistent downregulation across breeding stages. These genes point to a transcriptional state that preserves synaptic and structural readiness outside of the breeding season. For instance, TTR, which encodes transthyretin, maintains thyroid hormone transport, a pathway known to support adult neurogenesis and seasonal vocal plasticity (Orije et al., 2022; Raymaekers et al., 2017). Similarly, GRIA4—previously identified in the HVC (Lovell et al., 2008)—an AMPA receptor subunit, and CELSR3, an adhesion GPCR involved in dendritic arborization and axon guidance (Sigoillot et al., 2016), suggest that neuronal wiring are upregulated prior to reproductive activation. The presence of RPL6, a ribosomal protein, and PRMT1, a histone arginine methyltransferase, further indicate ongoing protein synthesis and epigenetic regulation (Gou et al., 2010; Nicholson et al., 2009). Together, these findings suggest that the nonbreeding HVC maintains a flexible transcriptional environment and is likely already establishing the neuronal connections important for breeding season song. These pathways subsequently become downregulated once breeding season starts.

A second group of genes was expressed at low levels in the nonbreeding period and showed consistent upregulation across the breeding season. NRCAM, a neuronal adhesion molecule, promotes axon guidance and synaptogenesis, supporting network reorganization (Sakurai, 2012). In parallel, OXR1 provides neuroprotection against oxidative stress, likely counteracting the metabolic costs of prolonged singing and reproductive activity (Oliver et al., 2011). These results suggest that breeding stages are marked by transcriptional programs that enhance neurotransmission, stabilize synaptic function, and protect against cellular stress.

A third cluster showed low expression in the nonbreeding season, elevated expression during pre-laying and decrease during incubation and then again increase in egg-laying. Genes in this group were strongly associated with cytoskeletal dynamics and fine-scale structural remodeling, suggesting that the onset of reproduction is characterized by intense synaptic and morphological reorganization. TTLL7, which modifies tubulin to regulate microtubule stability and transport, supports dendritic spine remodeling and axonal dynamics (Janke et al., 2005b; van Dijk et al., 2007). TENM4, a transmembrane protein required for axon guidance and synaptic specificity, likely contributes to the refinement of motor pathways critical for song performance (Gienapp et al., 2017). FNBP1, a regulator of actin cytoskeleton remodeling and vesicle endocytosis, further underlines the role of cytoskeletal plasticity in these stages (Lovell et al., 2008). The temporal profile of this cluster suggests that pre-laying and egg-laying involve peak levels of structural and synaptic reorganization, enabling the production of complex, high-performance songs most relevant during early reproductive interactions.

### Seasonal changes in hormones and song: TTR

Among the DEGs common to all non-breeding to breeding comparisons, TTR emerges as a particularly relevant gene due to its well-established role in TH transport and neuroplasticity (Orije et al., 2022). Although THs are well-known regulators of seasonal reproductive transitions via the hypothalamic–pituitary–gonadal axis (McNabb, 2007; Yoshimura, 2013), their role in song behavior and neural plasticity is just beginning to be explored (Orije et al., 2022; Raymaekers et al., 2017). We observed substantial upregulation of TTR in some individuals in the non-breeding season, followed by a marked decrease during the egg-laying, and a subsequent small increase during incubation. This pattern suggests a cyclic expression trajectory across the annual cycle, with TTR beginning to rise again as birds approach the transition back into the non-breeding state. Such dynamics are consistent with a role for TTR-mediated TH transport in preparing the system for seasonal transitions. In photoperiodic species, THs mediate the transition from photorefractoriness (a state of insensitivity to long days) to photostimulation, which triggers both gonadal growth and elevated testosterone concentration (McNabb, 2007; Orije et al., 2022; Yoshimura, 2013). By modulating local TH concentrations in the brain, TTR could serve as a molecular bridge linking broad physiological transitions to specifically neuroplasticity in the SCS, potentially helping to coordinate the timing of neural system readiness for later neurogenesis, vocal behavior, and reproductive state.

Our findings highlight TH as a potential upstream regulator involved in the seasonal onset of song, possibly through preparatory modulation of TH availability in the nonbreeding season (Sharma et al., 2020; D. Wang et al., 2022). TH has recently been implicated in promoting song learning and cognition (Orije et al., 2022), and TTR may contribute indirectly to these processes by facilitating thyroid hormone delivery to the brain and supporting neuroprotection (Liz et al., 2020; Mukai et al., 2009). While a direct role of TTR in regulating song behavior year-round remains to be demonstrated, its known functions suggest it could influence the neural substrates underlying song related plasticity. Therefore, TTR and TH may contribute to pre-breeding and early-breeding preparation of the SCS to promote neurogenesis, song-learning and/or singing activity at the onset of the breeding season (Lim et al., 2014). Given the known role of hormones such as testosterone on song quality and complexity in HVC, these findings suggest an additional broader seasonal neuroendocrine regulation of vocal and reproductive traits, where TH may act as an upstream molecular switch, initiating seasonal transitions in singing behavior, while testosterone fine-tunes its expression (Orije et al., 2020, 2022). While TH-driven reproductive transitions are well established, few studies have directly connected this endocrine signaling to SCS-specific gene expression, underscoring the novelty of our findings (exceptions, Ko et al., 2021, 2024; Orije et al., 2022).

We selected DEGs using a significance threshold of *p* < 0.01 together with logFC > |1|, but we also explored genes that met the *p* < 0.05 criterion without filtering by logFC, in order to identify other potentially relevant candidates. By doing so, we found two more candidates shared across the three comparisons: NCKAP1 and PARK7. NCKAP1 encodes a component complex involved in actin cytoskeleton remodeling, synaptic plasticity, and dendritic spine formation, processes that are fundamental for neural circuit reorganization (Guo et al., 2020; Noh et al., 2023). PARK7 functions as a multifunctional protein with roles in oxidative stress protection, mitochondrial homeostasis, and regulation of apoptosis (B. Wang et al., 2016; Zhang et al., 2021), it also has been implicated in sperm maturation and motility (Otčenášková et al., 2023), as well as in bone metabolism during reproductive stages (Zhang et al., 2021). Although these results should be interpreted with caution since they stem from a more exploratory analysis, they suggest additional connections between gene regulation in the HVC and reproductive as well as behavioral traits, which may be of interest for future studies.

### Seasonal neurogenesis and TTLL7

TTLL7 is a key gene that we identified in a co-expression module associated with seasonal increases in HVC volume. TTLL7 is a particularly interesting candidate gene due to its role in neuronal development through the regulation of microtubule dynamics, which are critical for axonal transport, cell division, and the maintenance of neuronal architecture (Janke et al., 2005a; Ping et al., 2023). Specifically, TTLL7 encodes a tubulin-glutamic acid ligase responsible for polyglutamylation, a post-translational modification essential for proper microtubule function (Genova et al., 2023). Polyglutamylation modulates interactions between microtubules and microtubule-associated proteins, influencing neuronal remodeling and synaptic stability. Disruptions in this process have been linked to neurodegenerative diseases and impaired neural plasticity (Janke et al., 2005b; van Dijk et al., 2007).

Given TTLL7’s role in microtubule modification, it likely contributes to the dynamic restructuring and neuron migration that occurs within HVC leading to the breeding season (Janke et al., 2005b; van Dijk et al., 2007). Microtubule dynamics facilitate axon and dendrite formation, intracellular transport, and overall neuronal morphology changes required for such plasticity. Interestingly, TTLL11, a paralog within the same family, directly interacts with doublecortin to promote polyglutamylation and microtubule formation (Sébastien et al., 2025). Doublecortin is widely recognized as a marker for neurogenesis, specifically used to identify new neuron recruitment within HVC in songbirds (Balthazart et al., 2008, 2016). Altogether, this underscores the potential involvement of TTLL7 in neurogenesis and neuronal restructuring of the SCS, highlighting its functional importance in seasonal neural plasticity.

### A testosterone-associated gene network

WGCNA analysis also revealed a potential testosterone-responsive gene network (Module 10). Module 10 was significantly associated with seasonal changes in testosterone. The function of the genes within the module potentially integrates chromatin regulation (via FAM169A), cytoskeletal dynamics (via CLASP1 and PPP1R12B), and signaling processes involving TMEM196, supporting seasonal neurogenesis and structural remodeling within the SCS. FAM169A, a nuclear envelope–associated protein, has been implicated in higher-order chromatin organization and could potentially contribute to hormone-driven transcriptional reprogramming in hormone-sensitive neurons (Maric et al., 2017; Roux et al., 2012; Vymetalkova et al., 2019). FAM169A and CLASP1—a microtubule-stabilizing protein involved in neurite outgrowth— upregulation in all breeding stages suggests a coordinated role in translating hormonal cues into structural plasticity (Kapitein et al., 2015; Maiato et al., 2003). PPP1R12B, while less studied in neural contexts, is associated with actomyosin regulation, might further contribute to cytoskeletal reorganization (Arimura et al., 2001; Fujioka et al., 1998). TMEM196, although not previously characterized in avian brains, shows detectable expression in neurons in mammalian cortex and basal ganglia (Zarrella et al., 2024). Its presence in a testosterone-correlated module suggests that it may represent a novel candidate gene contributing to hormone-sensitive plasticity and potentially to neurogenesis-related processes in the SCS. Taken together, the co-expression of these transcripts points toward a unified mechanism by which testosterone orchestrates cellular and molecular changes—including migration, chromatin remodeling, and circuit reorganization—to align behavior with reproductive status.

## Conclusions

Our findings underscore the potential critical interplay between gene expression, endocrine signaling, neural plasticity, and reproductive physiology in driving seasonal behavioral transitions. The identification of differentially expressed genes such as TTR and TTLL7, along with their correlation to key traits like plasma testosterone concentrations and HVC volume, provides valuable insight into the molecular mechanisms underlying seasonal neuroplasticity. TTR, TTLL7, and FAM169A may act at the intersection of endocrine signaling and neural remodeling. TTR, through its role in thyroid hormone transport, may also prime the SCS for seasonal transitions in song behavior, while TTLL7 may support neurogenesis via microtubule modification within HVC. Our co-expression analysis highlights genes in module 10 as a testosterone-responsive gene network, potentially integrating functions like chromatin regulation (FAM169A), cytoskeletal dynamics (CLASP1), and signaling processing (TMEM196). These findings support the idea that hormone-responsive gene networks coordinate chromatin and cytoskeletal regulation with additional signaling components to align neural plasticity with reproductive state. Further investigations are required to elucidate how these molecular mechanisms contribute to the structural and functional plasticity of the brain in species with seasonal reproductive patterns, which could offer new perspectives on the neuroendocrine processes governing neuronal plasticity in reproductive contexts.

