## Supplemental Material for "Seasonal neurogenomic changes provide genetic links between male song and testosterone-mediated neurogenesis in a wild songbird"

**Supplementary material**

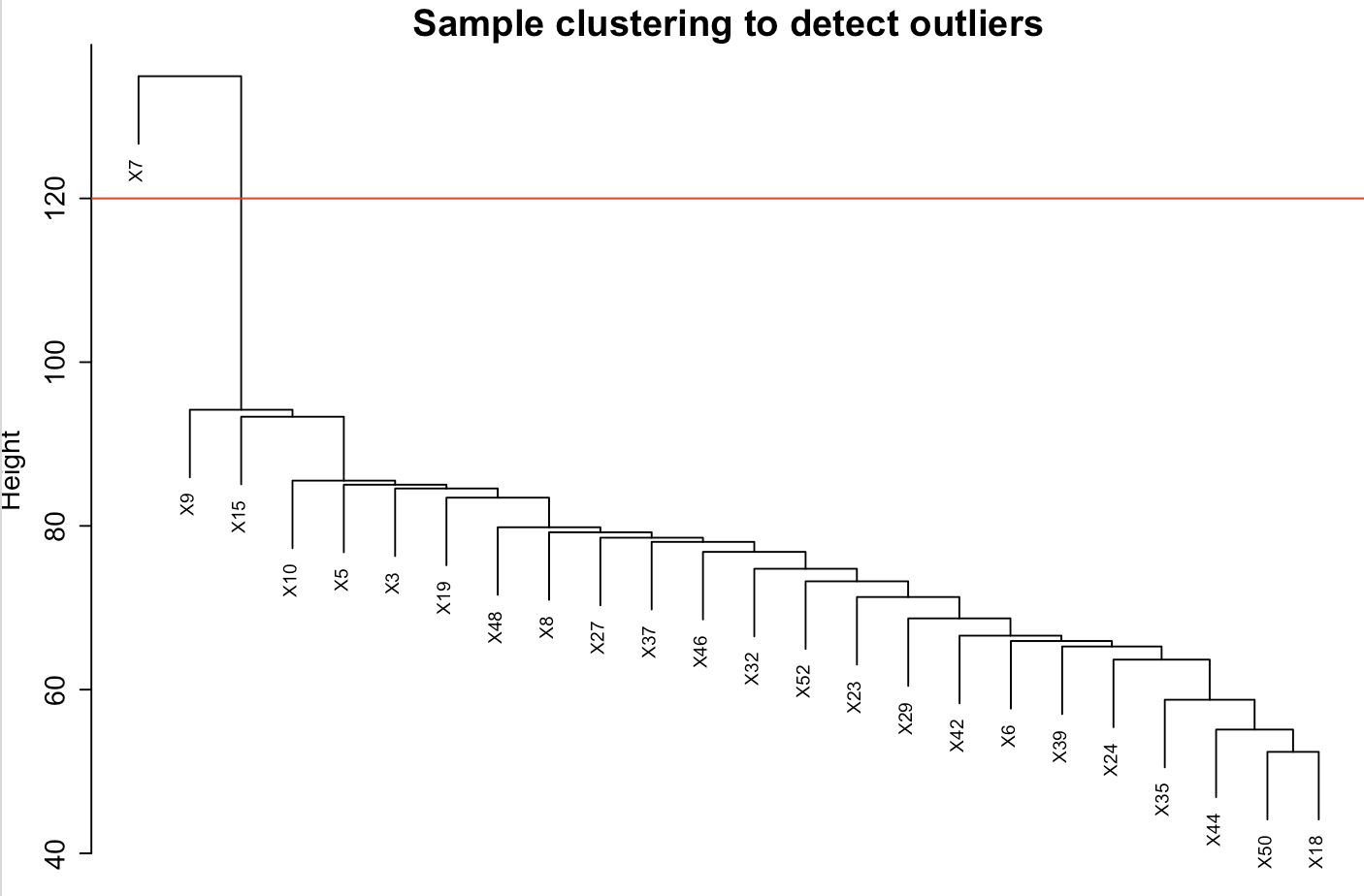

**Supplementary Figure 1.** Hierarchical clustering dendrogram to detect outlier samples in WGCNA analysis. The red horizontal line indicates the threshold for outlier detection. Samples that cluster above this threshold (e.g., X7) are identified as potential outliers and may be excluded from further analysis to ensure robust network construction.

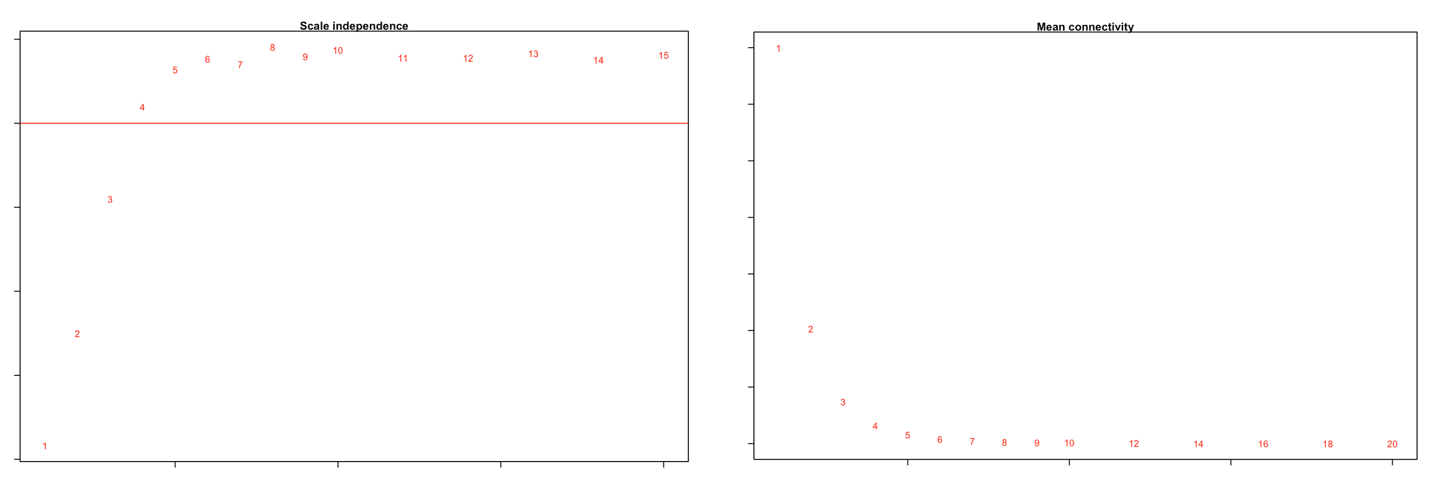

**Supplementary Figure 2. (a)** The scale independence plot displays the scale-free topology model fit (R²) as a function of the soft-thresholding power. The red horizontal line (R² = 0.90) indicates the threshold used to select an appropriate power, ensuring the network follows a scale-free topology**. (b)** The mean connectivity plot illustrates the mean gene connectivity as the soft-thresholding power increases. A decline in mean connectivity confirms the sparsity of the resulting network, consistent with the properties of a scale-free structure

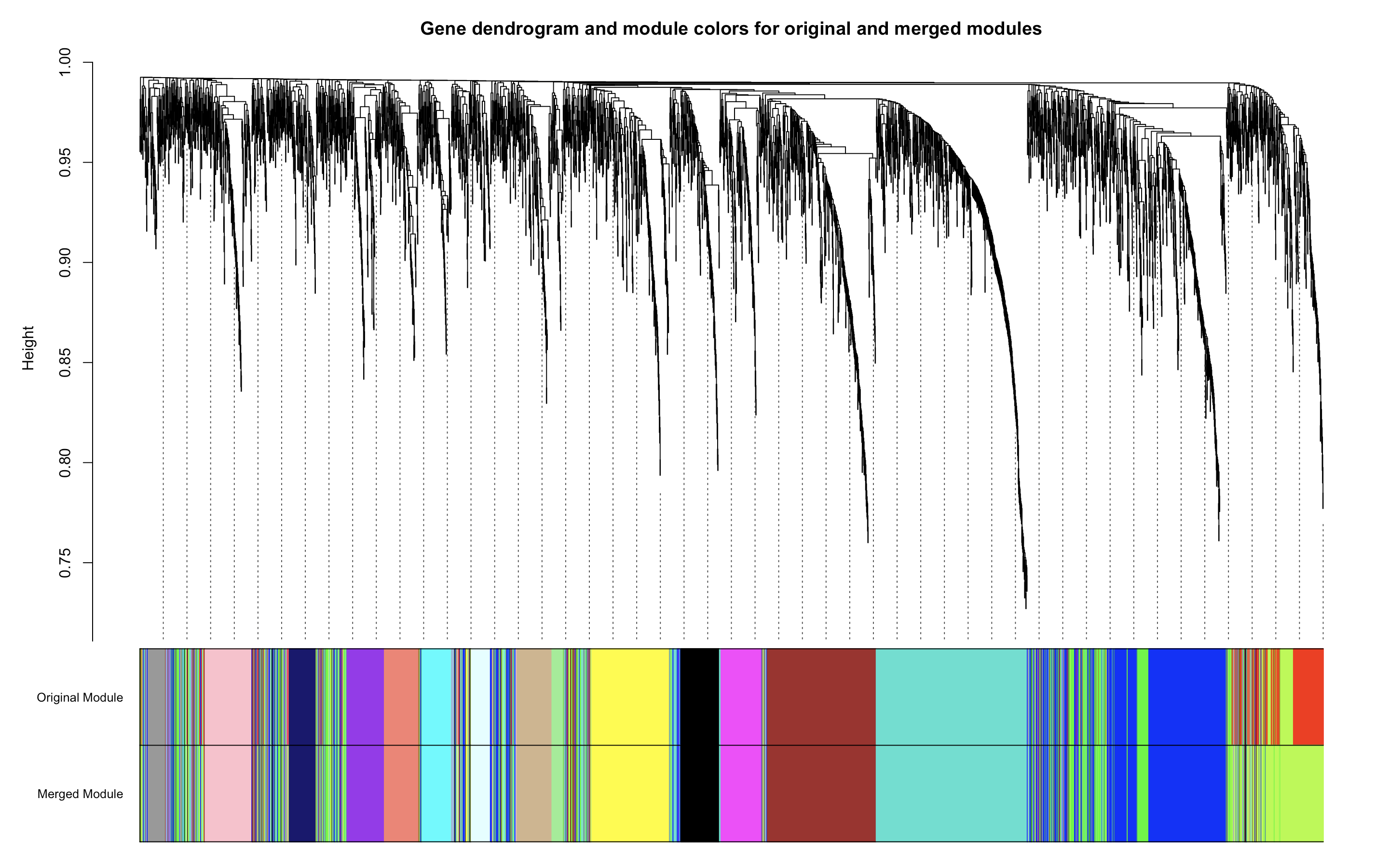

**Supplementary Figure 3.** Cluster dendrogram and module colors after module merging. The dendrogram represents the hierarchical clustering of genes based on their topological overlap, with branches corresponding to distinct gene modules. Initial module assignments are shown in the original colors, while merged modules after applying the module eigengene similarity threshold are displayed in the merged colors. The merging process reduces module redundancy while preserving biological relevance.

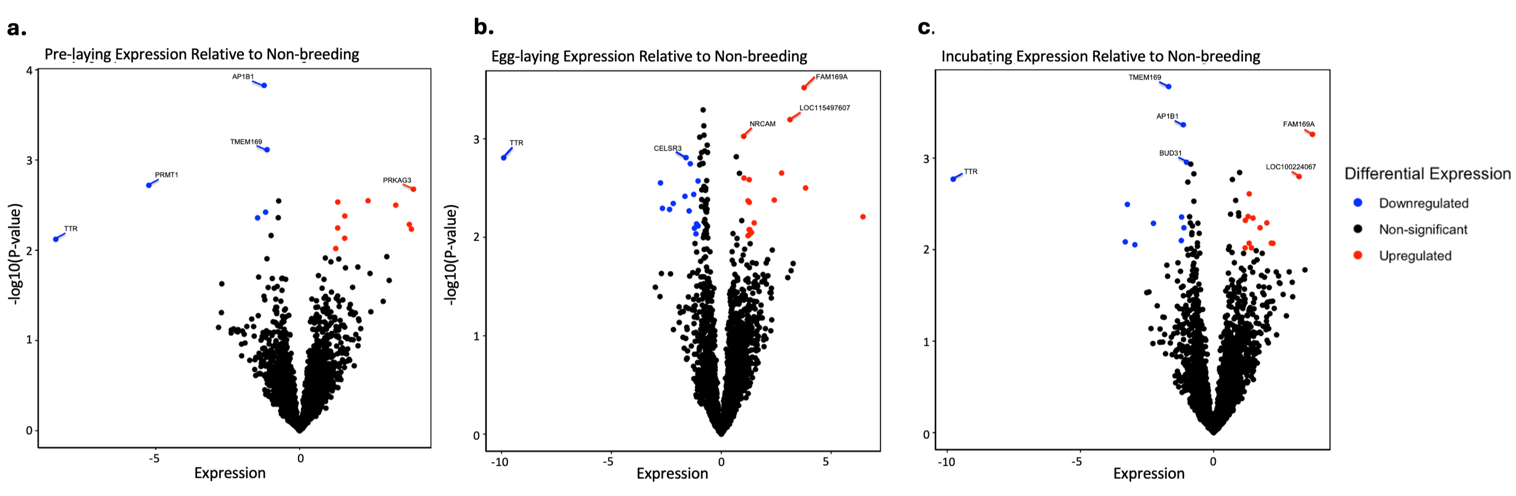

**Supplementary Figure 4**. Volcano plots comparing transcript expression during the non-breeding period with each seasonal stage (**a.** pre-laying, **b**. egg-laying, **c.** incubation). Transcripts with significant upregulation (log fold change > 1) are shown in red, while significantly downregulated transcripts (log fold change < -1) are shown in blue. The most differentially expressed transcript, TTR, stands out prominently.

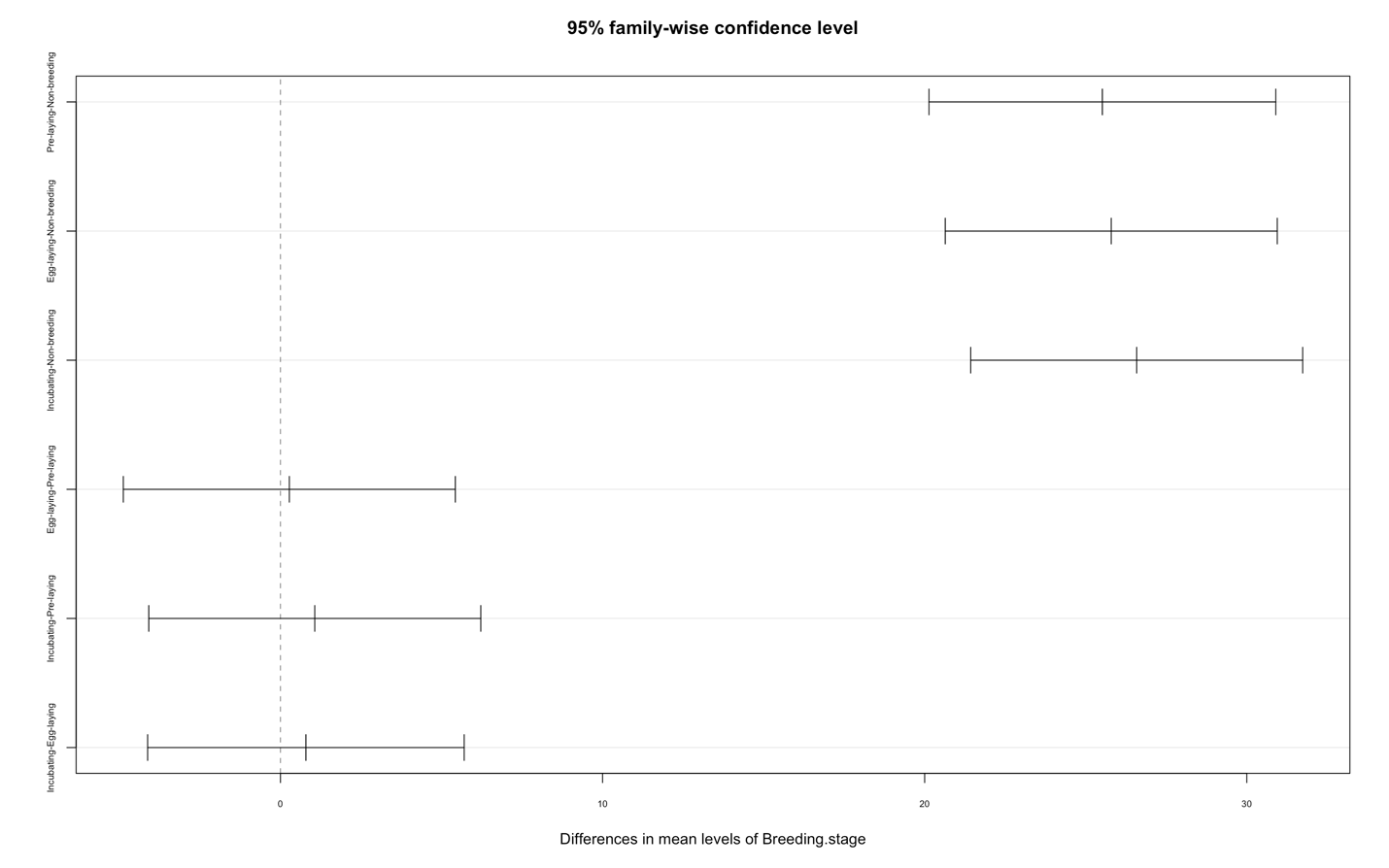

**Supplementary Figure 5.** Tukey's HSD plot for pairwise comparisons of testes size across breeding stages. The plot displays the mean differences between stages along with 95% confidence intervals. Significant differences are indicated where the confidence intervals do not overlap zero. The analysis reveals that testis size was significantly larger during the pre-laying, laying, and incubating stages compared to the non-breeding stage, with mean differences of 25.44 mm (p < 0.0001), 25.71 mm (p < 0.0001), and 26.50 mm (p < 0.0001), respectively.

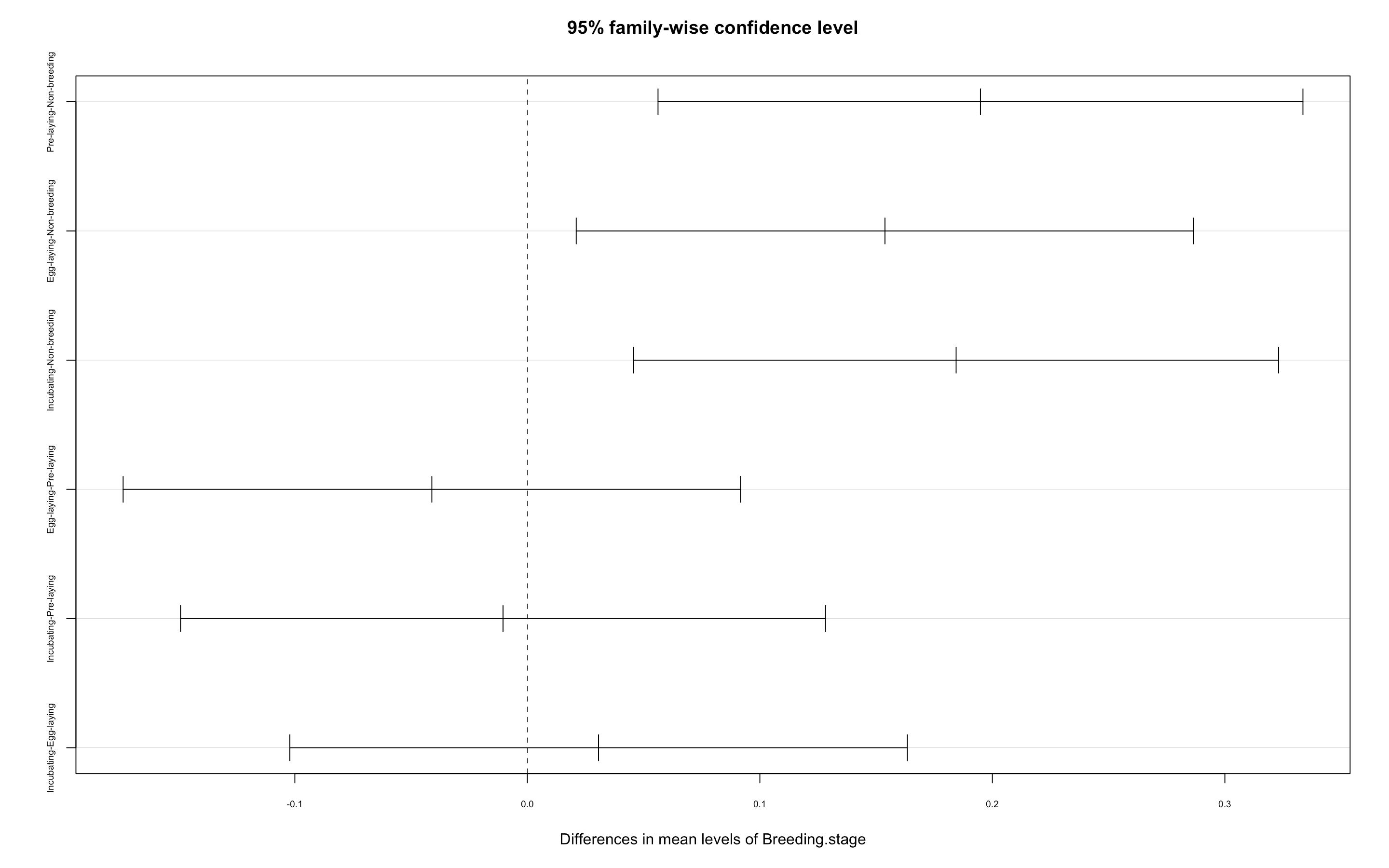

***Supplementary Figure 6.*** Tukey's HSD plot for pairwise comparisons of HVC volume (mm3) across breeding stages. The plot displays the mean differences between stages along with 95% confidence intervals. Significant differences are indicated where the confidence intervals do not overlap zero. The analysis reveals that testis size was significantly larger during the pre-laying, laying, and incubating stages compared to the non-breeding stage, with mean differences of 0.19 mm (p = 0.004), 0.153 mm (p = 0.020), and 0.184 mm (p = 0.007), respectively.

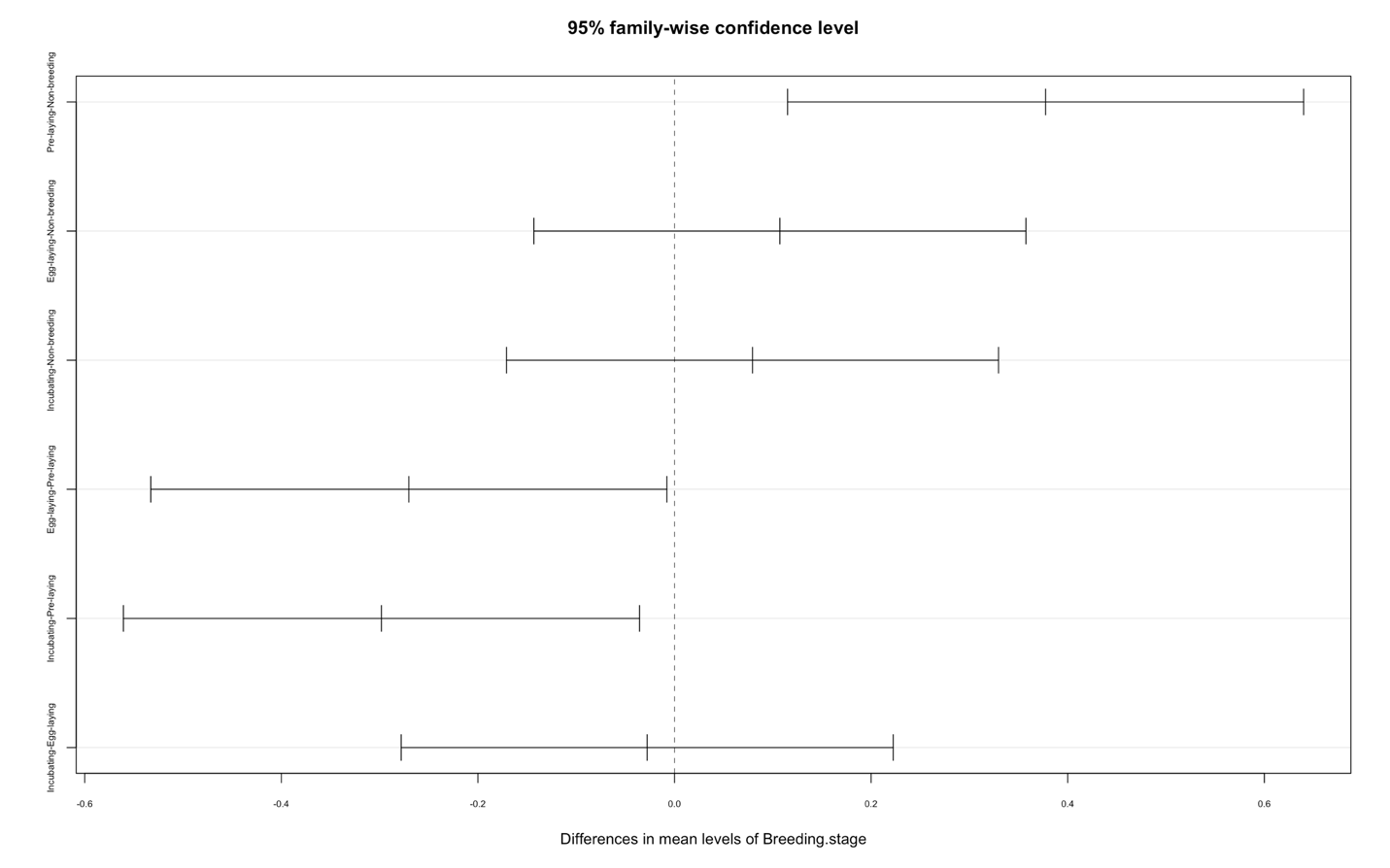

***Supplementary Figure 7.*** Tukey's HSD plot for pairwise comparisons of testosterone concentration (ng/mL) across breeding stages. The plot displays the mean differences between stages along with 95% confidence intervals. Significant differences are indicated where the confidence intervals do not overlap zero. The analysis reveals a significant difference between the pre-laying and non-breeding stage (mean difference = 0.377 ng/mL, p = 0.003), egg-laying and pre-laying (mean difference = -0.270ng/mL, p = 0.042) and incubating and pre-laying (mean difference = -0.298ng/mL, p = 0.022).

**Supplementary Table 1:** sample information

| **Bird ID** | | **Sex** | **Breeding stage** | | **Date** | | **Time** | | **County** | | **State** | | **Location details (SGL = State Game Land** |
| --- | --- | --- | --- | --- | --- | --- | --- | --- | --- | --- | --- | --- | --- |
| HOWR3 | M | Non-breeding | | 2/14/2022 | | 9:15 | | Alachua | | FL | | University of Florida, Hague Dairy | |
| HOWR5 | M | Non-breeding | | 2/15/2022 | | 8:35 | | Alachua | | FL | | University of Florida, Hague Dairy | |
| HOWR6 | M | Non-breeding | | 2/15/2022 | | 16:00 | | Alachua | | FL | | University of Florida, Hague Dairy | |
| HOWR7 | M | Non-breeding | | 2/16/2022 | | 8:15 | | Alachua | | FL | | University of Florida, Hague Dairy | |
| HOWR8 | M | Non-breeding | | 2/17/2022 | | 9:48 | | Alachua | | FL | | University of Florida, Hague Dairy | |
| HOWR9 | M | Non-breeding | | 2/17/2022 | | 12:50 | | Alachua | | FL | | University of Florida, Hague Dairy | |
| HOWR10 | M | Non-breeding | | 2/17/2022 | | 17:30 | | Alachua | | FL | | University of Florida, Hague Dairy | |
| HOWR15 | M | Pre-laying | | 5/8/2022 | | 12:05 | | Wanye | | PA | | SGL310.20 | |
| HOWR18 | M | Pre-laying | | 5/9/2022 | | 8:10 | | Wanye | | PA | | SGL310.25 | |
| HOWR19 | M | Pre-laying | | 5/9/2022 | | 17:35 | | Wanye | | PA | | SGL310.10 | |
| HOWR23 | M | Egg-laying | | 5/30/2022 | | 9:50 | | Wanye | | PA | | SGL310.25 | |
| HOWR24 | M | Incubation | | 7/8/2022 | | 5:50 | | Lackawanna | | PA | | SGL300 | |
| HOWR27 | M | Incubation | | 7/9/2022 | | 6:30 | | Wanye | | PA | | SGL310.1 | |
| HOWR29 | M | Pre-laying | | 5/11/2023 | | 6:30 | | Lackawanna | | PA | | SGL300 | |
| HOWR32 | M | Pre-laying | | 5/11/2023 | | 14:45 | | Wanye | | PA | | SGL310.25 | |
| HOWR35 | M | Egg-laying | | 5/18/2023 | | 6:00 | | Lackawanna | | PA | | SGL300 | |
| HOWR37 | M | Egg-laying | | 5/19/2023 | | 6:05 | | Wanye | | PA | | SGL310.10 | |
| HOWR39 | M | Egg-laying | | 5/20/2023 | | 5:40 | | Wanye | | PA | | SGL310.25 | |
| HOWR42 | M | Egg-laying | | 5/25/2023 | | 5:38 | | Wanye | | PA | | SGL310.1 | |
| HOWR44 | M | Incubation | | 5/29/2023 | | 5:50 | | Lackawanna | | PA | | SGL300 | |
| HOWR46 | M | Incubation | | 5/30/2023 | | 6:30 | | Wanye | | PA | | SGL310.1 | |
| HOWR48 | M | Incubation | | 5/31/2023 | | 5:45 | | Wanye | | PA | | SGL310.10 | |
| HOWR50 | M | Egg-laying | | 6/2/2023 | | 6:06 | | Lackawanna | | PA | | SGL300 | |
| HOWR52 | M | Incubation | | 6/4/2023 | | 6:42 | | Wanye | | PA | | SGL310.25 | |

**Supplementary Table 2:** Highest percentage of DEGs relative to the total number of genes in each module

| **Module** | **% of DEGs relative to the total number of genes** |
| --- | --- |
| 0 | 0.0068 |
| 1 | 0.0336 |
| 2 | 0.0140 |
| 3 | 0.0152 |
| 4 | 0.0117 |
| 5 | 0.0087 |
| 6 | 0.0100 |
| 7 | 0.0120 |
| 8 | 0.0127 |
| 9 | 0.0137 |
| 10 | 0.0323 |
| 11 | 0.0000 |
| 12 | 0.0175 |
| 13 | 0.0187 |
| 14 | 0.0000 |
| 15 | 0.0227 |
| 16 | 0.0000 |
| 17 | 0.0000 |
| 18 | 0.0238 |
